# Contrasting effects of global and local cortical activity on the regulation of sleep

**DOI:** 10.1101/2025.09.07.673922

**Authors:** Lukas B. Krone, Jack D. Hamilton, Anna Hoerder-Suabedissen, Clara Marnitz, Lale Memis, Anna B. Szabo, Colin J. Akerman, Zoltán Molnár, Vladyslav V. Vyazovskiy

## Abstract

The cortex is now recognised as a key regulator of sleep. It remains unclear, however, how sleep is affected by changes in cortical activity over different spatial scales. For instance, sleep states are associated with ‘global’ changes in activity across cortex, but sleep pressure is associated with ‘local’ variations in cortical activity. Here we use chemogenetic manipulations of a single neuronal population to determine how sleep reflects the spatial scale of cortical activity. Global inhibition of *Rbp4-Cre+* neurons increases sleep, but also produces abnormal brief-awakenings and reduces electroencephalogram slow-wave activity, a marker of sleep intensity. In contrast, local inhibition/excitation of *Rbp4-Cre+* neurons in the prefrontal but not occipital cortex increases sleep/wake, respectively, whilst preserving normal sleep architecture and electrophysiological features. We conclude that globally altering cortical activity may induce unphysiological sleep due to conflicting sleep-regulatory signals, while locally modulating prefrontal cortex affords bidirectional sleep regulation that leaves naturalistic features intact.

## Introduction

Sleep has been quantified for almost a century using readouts from the cerebral cortex, such as the amplitude of slow waves in the electroencephalogram (EEG)^1^, and slow-wave activity (SWA, 0.5-4 Hz) during non-rapid eye movement (NREM) sleep^2,3^. The cerebral cortex itself is strongly influenced by sleep-regulatory substances^4^ and harbours neuronal populations that are sleep-active and reflect the level of homeostatic sleep drive^5,6^. Yet, the idea that the cerebral cortex actively contributes to the regulation and control of sleep is a recent one^7–9^.

We previously demonstrated an active role for the cerebral cortex in sleep regulation using mice where *Rbp4-Cre+* layer 5 pyramidal neurons and dentate gyrus granule cells were functionally silenced across the cortex^10^. These mice displayed a three-hour reduction in daily sleep time and a suppressed homeostatic sleep response to extended wakefulness^10^. This is in line with the idea that the phenomenon of sleep is tightly coupled to ‘global’ cortical activity and that, as a behaviour, sleep usually affects the whole animal.

Interestingly, sleep pressure is thought to accumulate on a ‘local’ level of individual brain cells and cortical columns^11,12^, and ‘local sleep’ (i.e. localised sleep-like activity patterns in the neocortex) can even occur during extended wakefulness^13^. The prefrontal cortex (PFC) appears to have a particular relevance for local and global sleep. For example, it has long been postulated that the PFC ‘gets tired first’^14^ based on the frontocentral pattern of EEG synchronisation during extended wakefulness and sleep onset^15^, the fronto-posterior gradient in the level of SWA during NREM sleep^16–18^, and the typical frontocentral origin of slow waves^19,20^.

Recent work demonstrates that the PFC harbours important sleep regulatory neurons. Stimulating PFC*^Sst^*^-GABA^ neurons can elicit sleep preparation and initiation^21^; excitatory neurons in the medial PFC can initiate and maintain rapid eye movement (REM) sleep^22^; tetrodotoxin-inactivation of the mPFC suppresses REM sleep and increases NREM sleep^23^; and chemogenetically increasing synaptic spine size of pyramidal neurons of the PFC increases sleep NREM amount^24^. Interestingly, other cortical regions contribute different aspects of sleep regulation. For example, excitatory neurons in the retrosplenial and primary visual cortices have an influence over REM sleep amount, features, and sub-states^25,26^. Changes in the activity of specific ‘local’ subpopulations of cortical neurons - defined by their cortical region, layer, and/or molecular entities - might therefore explain their distinct sleep-regulatory properties. How this compares to the role of global cortical activity in determining sleep amount, intensity, and characteristics remains unclear.

To investigate this, we tested how the activity of a distinct cortical neuronal population modulated over different spatial scales – that is, globally versus locally – might elicit differential effects on sleep. To achieve this, we utilized different expression strategies for Designer Receptors Exclusively Activated by Designer Drugs (DREADDs), modulating the activity of *Rbp4-Cre+* layer 5 pyramidal neurons globally across the cortex via transgenics, or locally in the prefrontal and occipital cortices via viral vectors. In each case we acutely altered *Rbp4-Cre+* neuronal activity via administration of clozapine-N-oxide (CNO) and observed the effects on subsequent sleep using EEG and electromyogram (EMG) recordings.

We show that acute inhibition of *Rbp4-Cre+* neurons on a global cortical scale increases sleep amount, but that this sleep is fragmented and of reduced intensity. In contrast, local modulations of *Rbp4-Cre+* neuronal activity in the PFC alone can bidirectionally alter sleep amount but leave naturalistic sleep features intact. Local *Rbp4-Cre+* modulations in the occipital cortex (OCC) bidirectionally alter several architectural parameters but not sleep amount. Together, our data demonstrate that targeted inhibition/excitation of *Rbp4-Cre+* neurons in the PFC can induce/suppress physiological sleep, respectively, but that this cannot be achieved via the same neuronal population when affected globally or in the OCC. We shed light on how different spatial scales of acute cortical activity have contrasting effects on sleep, and provide a local target in the PFC by which sleep can be effectively and physiologically modulated.

## Results

### ‘Global’ cortex-wide chemogenetic inhibition of *Rbp4-Cre+* neurons promotes sleep

To investigate how a widespread (‘global’) manipulation of cortical *Rbp4-Cre+* neurons acutely impacts sleep, we first crossed the pan-layer 5 driver line Rbp4^Cre^ to the inhibitory DREADD-reporter line R26-LSL-Gi-DREADD (R26^LSL-hM4Di^) and performed chemogenetic manipulations combined with chronic EEG/EMG recordings in young adult male Rbp4^hM4Di^ mice (Fig. 1a,b).

**Figure 1:**
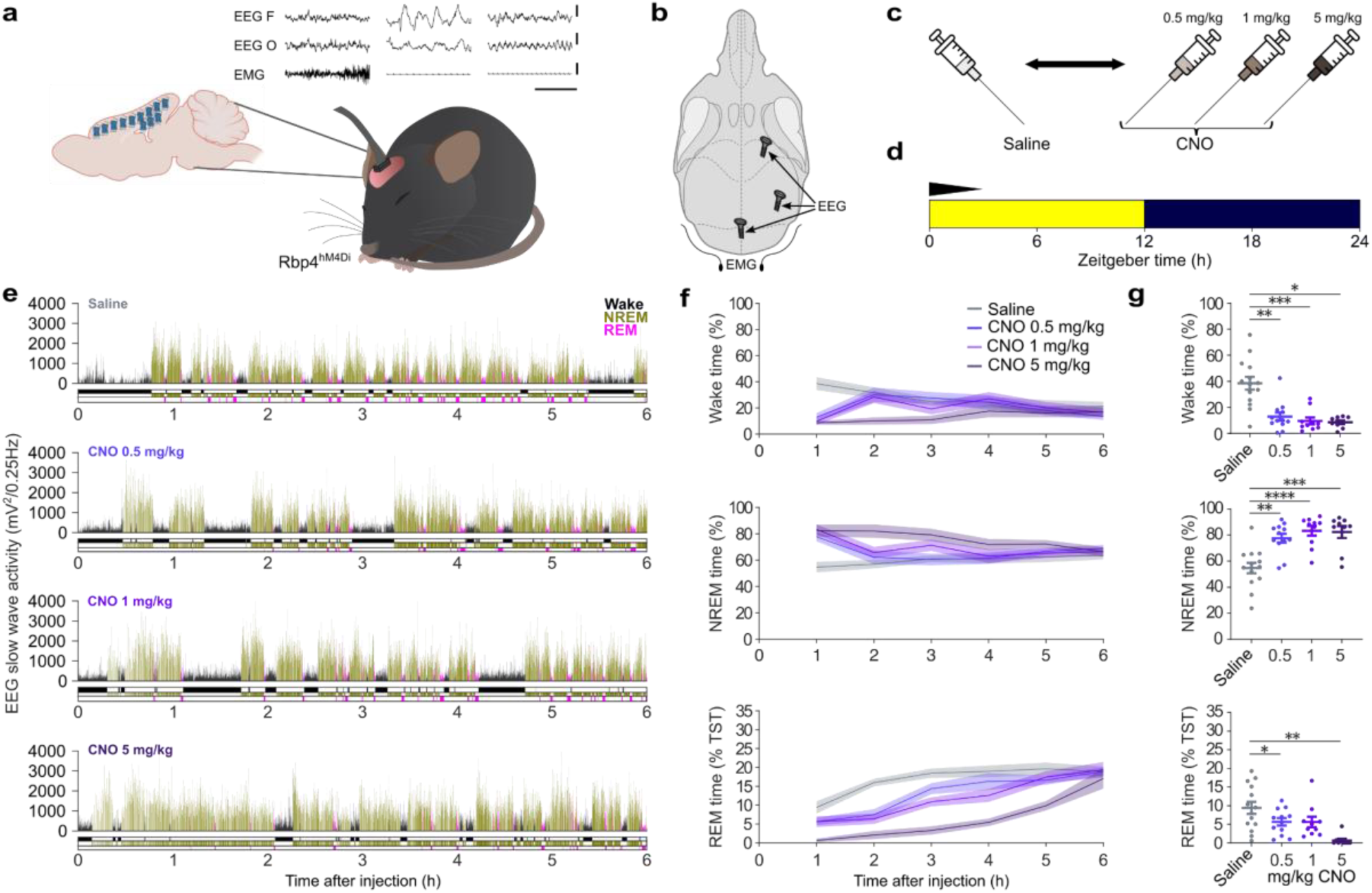
Cortex-wide chemogenetic inhibition of *Rbp4-Cre+* neurons increases sleep. **(a)** Schematic of electrophysiological recordings in Rbp4^hM4Di^ mice. Scale bars represent 500 μV on the y-axis and 1 s on the x-axis. **(b)** Schematic of electroencephalogram (EEG) recording electrodes in the skull. **(c and d)** Experimental paradigm: counterbalanced intraperitoneal (i.p.) injections of saline versus 0.5 mg/kg, 1 mg/kg, and 5.0 mg/kg clozapine-*N*-oxide (CNO) were administered at light onset (ZT=0). **(e)** Representative hypnograms and EEG slow-wave activity (SWA, 0.5-4 Hz, 4-s epochs) of a mouse following injections of saline (top panel) and the three CNO doses (bottom three panels). **(f)** Time courses of Wake, NREM, and REM sleep in the first 6 hr post-i.p. injection of saline and each of the three CNO doses, with **(g)** corresponding quantifications of time spent in each vigilance state during the first hour of DREADD activation (saline = grey; light purple = 0.5 mg/kg CNO; medium purple = 1.0 mg/kg CNO; dark purple = 5.0 mg/kg CNO). Rapid eye movement (REM) sleep is presented as a percentage of total sleep time (% TST). Note that all time analyses were aligned from 15 min post-injection (as T0) to ensure full DREADD activation and are visualised as mean ± SEM (shaded). Asterisks indicate comparisons with significant differences (*p<0.05; **p<0.01; ***p<0.001, ****p<0.0001) for analyses with significant main effects. n = 14, 12, 10, 9 for saline, 0.5 mg/kg, 1.0 mg/kg, 5 mg/kg CNO, respectively, for panels f and g. CNO: Clozapine-*N*-oxide; DREADD: Designer Receptors Exclusively Activated by Designer Drugs; EEG: Electroencephalogram; EEG F: Frontal EEG; EEG O: Occipital EEG; EMG: Electromyography; i.p.: intraperitoneal; NREM: Non-rapid eye movement sleep; REM: Rapid eye movement sleep; SWA: Slow wave activity; TST: Total sleep time; T0: timepoint of analysis onset 15min post-injection; ZT: Zeitgeber time.

In pilot experiments to optimise CNO dosage, we injected mice intraperitoneally (i.p.) with three doses of CNO (0.5 mg/kg, 1 mg/kg, 5 mg/kg) at light onset (Fig. 1c,d). In the first hour of DREADD activation (0:15 - 1:15h after i.p. injections), all three CNO doses elicited a significant increase in NREM sleep time (saline: 54.7±14.92%, CNO 0.5 mg/kg: 77.64±11.7%, CNO 1 mg/kg: 83.31±11.77%, CNO 5 mg/kg: 82.36±13.68%; condition effect: *F*(2.645, 24.69) = 19.08, *p* < 0.0001); and reduction in Wake time (saline: 38.56±18.36%, CNO 0.5 mg/kg: 12.97±11.02%, CNO 1 mg/kg: 9.67±8.43%, CNO 5 mg/kg: 8.62±4.26%; dose effect: *F*(1.669, 15.58) = 19.41, *p* = 0.002) compared to saline (CNO 0.5 mg/kg: *p* = 0.002, CNO 1 mg/kg: p < 0.0001, CNO 5 mg/kg: *p* = 0.001 for NREM and CNO 0.5 mg/kg: *p* = 0.002, CNO 1 mg/kg: *p* = 0.001, CNO 5 mg/kg: p = 0.012 for wakefulness post-hoc comparisons, respectively) (Fig. 1f,g). Beyond the first hour, only the high CNO dose (5 mg/kg) elicited a sustained effect, increasing NREM sleep and reducing Wake compared to saline for approximately 3-4 hours (Fig. 1f). In fact, the 5 mg/kg CNO dose caused such a profound response that shortly after injection, mice began engaging in sleep preparatory behaviours and then fell asleep, even when exposed to a novel object which usually stimulates mice to stay awake ([Suppl. Video 1](https://figshare.com/articles/media/Supplementary_Video_1/30069778)).

The amount of REM sleep as a percentage of total sleep time (% TST) was reduced by all three CNO doses (CNO 0.5 mg/kg: 5.65±3.47% TST, CNO 1 mg/kg: 5.60±4.46% TST, CNO 5 mg/kg: 0.63±1.45% TST) compared to saline (saline: 9.39±6.08% TST) (Fig. 1f,g). The 0.5 mg/kg and 5 mg/kg doses yielded statistically significant effects in the first hour post-DREADD activation (dose effect: F(1.786, 16.67) = 7.847, *p* = 0.005; post hoc: CNO 0.5 mg/kg: *p* = 0.045, CNO 1 mg/kg: p = 0.288, CNO 5 mg/kg: *p* = 0.009 for wakefulness post-hoc comparisons) (Fig. 1g).

Since the lowest dose of CNO (0.5 mg/kg) could reliably increase the amount of NREM sleep (Fig. 1e,f,g), we conducted all subsequent experiments at this dosage, counterbalancing injections of 0.5 mg/kg CNO and saline. Sleep and wake amounts remained unchanged in DREADD-free R26^LSL-hM4Di^ controls injected with this CNO dose (Suppl. Fig. 1).

### ‘Local’ prefrontal chemogenetic inhibition and excitation of *Rbp4-Cre+* neurons bidirectionally changes sleep amount

Given that sleep starts on a ‘local’ level, including regions of cortex and populations of cortical neurons^11,13^, we next asked whether chemogenetic inhibition of *Rbp4-Cre+* layer 5 pyramidal neurons in the PFC alone could elicit an acute sleep effect. Our choice of the PFC was informed by the known role of centrofrontal cortical regions in the initiation of slow waves^19^ and sleep behaviours^21^. Here, we injected adeno-associated virus for Cre-dependent expression of inhibitory (*pAAV8-hSyn-DIO-hM4D(Gi)-mCherry*) DREADD receptors in the PFC of Rbp4^Cre^ mice (Fig. 2a,b).

**Figure 2:**
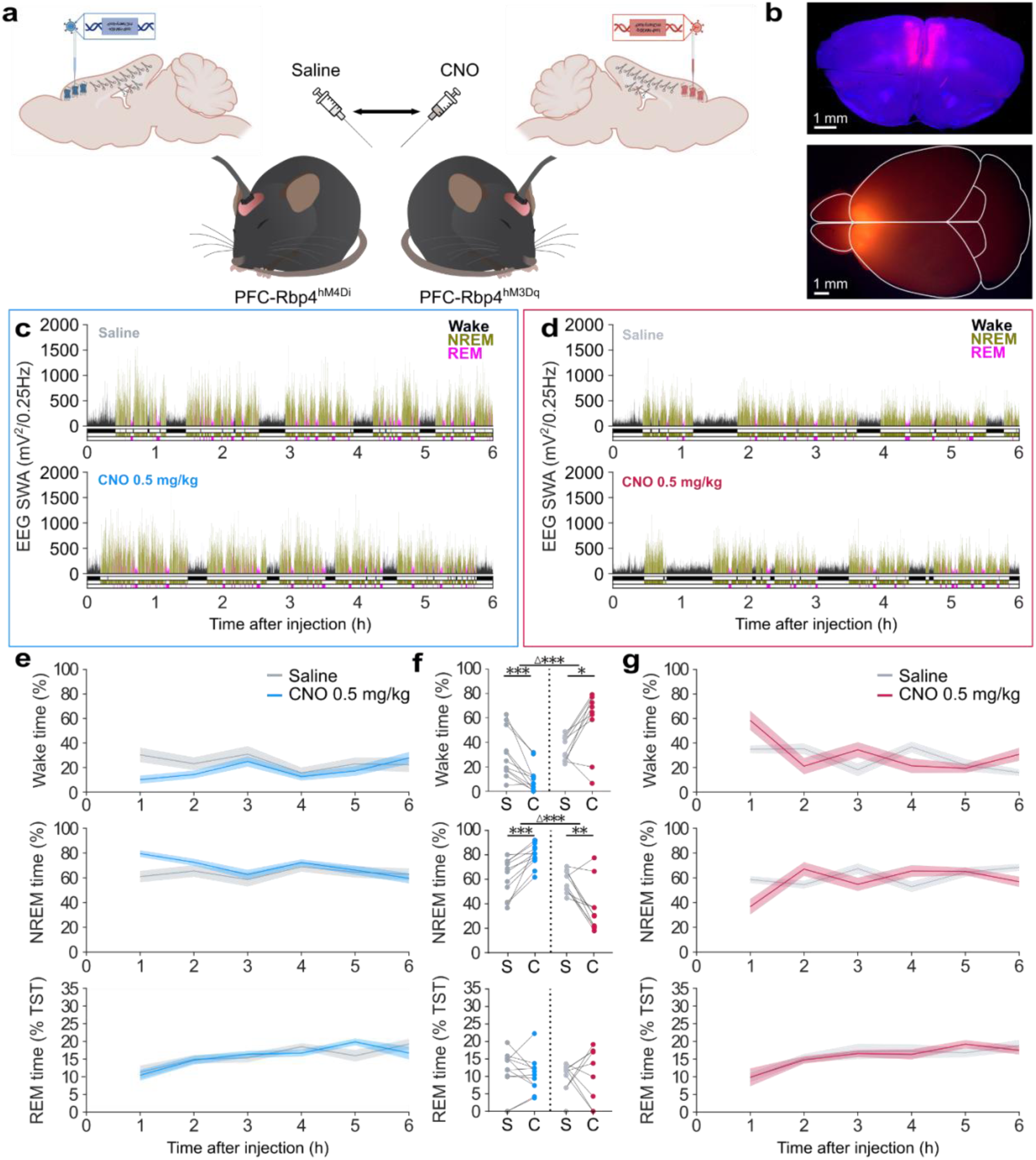
Targeted, chemogenetic inhibition/excitation of prefrontal cortical layer 5 pyramidal neurons bidirectionally alter sleep amount. **(a)** Schematic of viral vector injections (inhibitory hM4Di, cyan; excitatory hM3Dq, red) for localised DREADD expression in the prefrontal cortex (PFC) of Rbp4^Cre^ mice. Note Cre expression is represented as biological scissors. Experiments consisted of counterbalanced intraperitoneal (i.p.) injections of saline vs. 0.5 mg/kg CNO at light onset (ZT=0). **(b)** Coronal brain section (top) and dissection microscope image (bottom) showing PFC DREADD expression. Scale bars, 1mm. **(c and d)** Representative hypnograms and EEG slow-wave activity (SWA, 0.5-4 Hz, 4-s epochs) of **(c)** PFC-Rbp4^hM4Di^ and **(d)** PFC-Rbp4^hM3Dq^ mice following i.p. injections of saline (top) and 0.5 mg/kg clozapine-*N*-oxide (CNO) (bottom) at ZT=0. **(e and g)** Time courses of Wake, NREM, and REM sleep in the first 6 hr post-injection of saline and 0.5 mg/kg CNO in **(e)** PFC-Rbp4^hM4Di^ and **(g)** PFC- Rbp4^hM3Dq^ mice. **(f)** Percentages of time spent in Wake, NREM, and REM sleep during the first hour of DREADD activation in PFC-Rbp4^hM4Di^ (cyan) and PFC-Rbp4^hM3Dq^ (red) mice, compared to saline (grey, S). Rapid eye movement (REM) sleep is presented as a percentage of total sleep time (% TST). Note that all time analyses were aligned from 15 min post-injection (as T0) to ensure full DREADD activation and are visualised as mean ± SEM (shaded). Asterisks indicate comparisons with significant differences (*p<0.05; ***p<0.001, ****p<0.0001;) for analyses with significant main effects. n=11 PFC-Rbp4^hM4Di^ for panel e, n=10 PFC-Rbp4^hM3Dq^ for panel g, n=11 PFC-Rbp4^hM4Di^ and n=10 PFC-Rbp4^hM3Dq^ for panel f. C: CNO; CNO: Clozapine-*N*-oxide; DREADD: Designer Receptors Exclusively Activated by Designer Drugs; EEG: Electroencephalogram; EMG: Electromyogram; i.p.: intraperitoneal; NREM: Non-rapid eye movement sleep; PFC: Prefrontal cortex; REM: Rapid eye movement sleep; S: Saline; SWA: Slow-wave activity; TST: Total sleep time; T0: timepoint of analysis onset 15min post-injection; ZT: Zeitgeber time.

Activation of DREADDs in PFC-Rbp4^hM4Di^ mice significantly increased overall sleep amount (Fig. 2c,e,f) to a similar extent as in the global Rbp4^hM4Di^ mice, despite inhibiting a substantially smaller number of *Rbp4-Cre+* neurons. Specifically, the amount of NREM sleep in PFC-Rbp4^hM4Di^ mice was significantly increased during the first hour of the DREADD manipulation compared to saline (saline: 60.98±15.32%, CNO 0.5 mg/kg: 79.52±9.03%; *t*(10) = 5.113, *p* = 0.001) (Fig. 2e,f), while Wake was significantly reduced (saline: 30.32±19.71%, CNO 0.5 mg/kg: 10.16±11.1%; *t*(10) = 4.654, *p* = 0.001) (Fig. 2e,f). REM sleep amount was unchanged (saline: 11.29±6.25% TST, CNO 0.5 mg/kg: 10.43±5.11% TST; *t*(10) = 0.578, *p* = 0.576) (Fig. 2e,f).

To test whether sleep could be bidirectionally modulated via PFC-*Rbp4-Cre+* neurons, we also injected the excitatory DREADD virus (*pAAV8-hSyn-DIO-hM3D(Gq)-mCherry)* in the PFC of Rbp4^Cre^ mice (Fig. 2a,b). In direct opposition to PFC-Rbp4^hM4Di^ mice, PFC-Rbp4^hM3Dq^ mice had significantly increased Wake time (Mdn_saline_ = 35.06%, IQR_saline_ = 19.11%, Mdn_CNO_ = 66.56%, IQR_CNO_ = 28.14%; *W* = 45, *p* = 0.02) (Fig. 2f,g) and reduced NREM sleep (saline: 58.56±8.734%, CNO 0.5 mg/kg: 36.66±20.04%; *t*(9) = 3.675, *p* = 0.005) (Fig. 2f,g) in the first hour of DREADD manipulations compared to saline (Fig. 2d,f,g). Like PFC-Rbp4^hM4Di^ mice, REM sleep amount was unaffected (Mdn_saline_ = 11.76%, IQR_saline_ = 5.67%, Mdn_CNO_ = 11.81%, IQR_CNO_ = 17.2%; *W* = 1, *p* > 0.999) (Fig. 2f,g).

To quantify the observed bidirectionality, we additionally assessed between-group differences, calculating the within-subject difference between CNO and saline conditions. We observed significant inhibitory versus excitatory effects for Wake time (Δ inhibitory: −20.16±14.37%, Δ excitatory: +23.51±23.10%; *t*(14.8) = 5.142, *p* = 0.0001) and NREM sleep (Mdn_Δ inhibitory_ = +17.44%, IQR_Δ inhibitory_ = 9.34%, Mdn_Δ excitatory_ = −30.39%, IQR_Δ excitatory_ = 27.81%; *U* = 5, *p* = 0.0001) (Fig. 2f), while no difference was observed for REM sleep amount (Δ inhibitory: −0.86±4.95%, Δ excitatory: +0.05±8.56%; *t*(14.13) = 0.296, *p* = 0.772) (Fig. 2f). Saline conditions were comparable between groups.

### ‘Local’ prefrontal but not ‘global’ cortical inhibition of *Rbp4-Cre+* neurons preserves physiological sleep features

Whilst global and local inhibition of cortical activity produced a similar increase in the total amount of NREM sleep, this does not mean that the sleep elicited was equivalent. For instance, sleep can vary in terms of how consolidated or uninterrupted it is, and how effectively it dissipates sleep pressure. To investigate whether the globally- and locally-induced sleep in our experiments differed, we directly contrasted the physiological sleep features of Rbp4^hM4Di^ and PFC-Rbp4^hM4Di^ mice (Fig. 3a).

**Figure 3:**
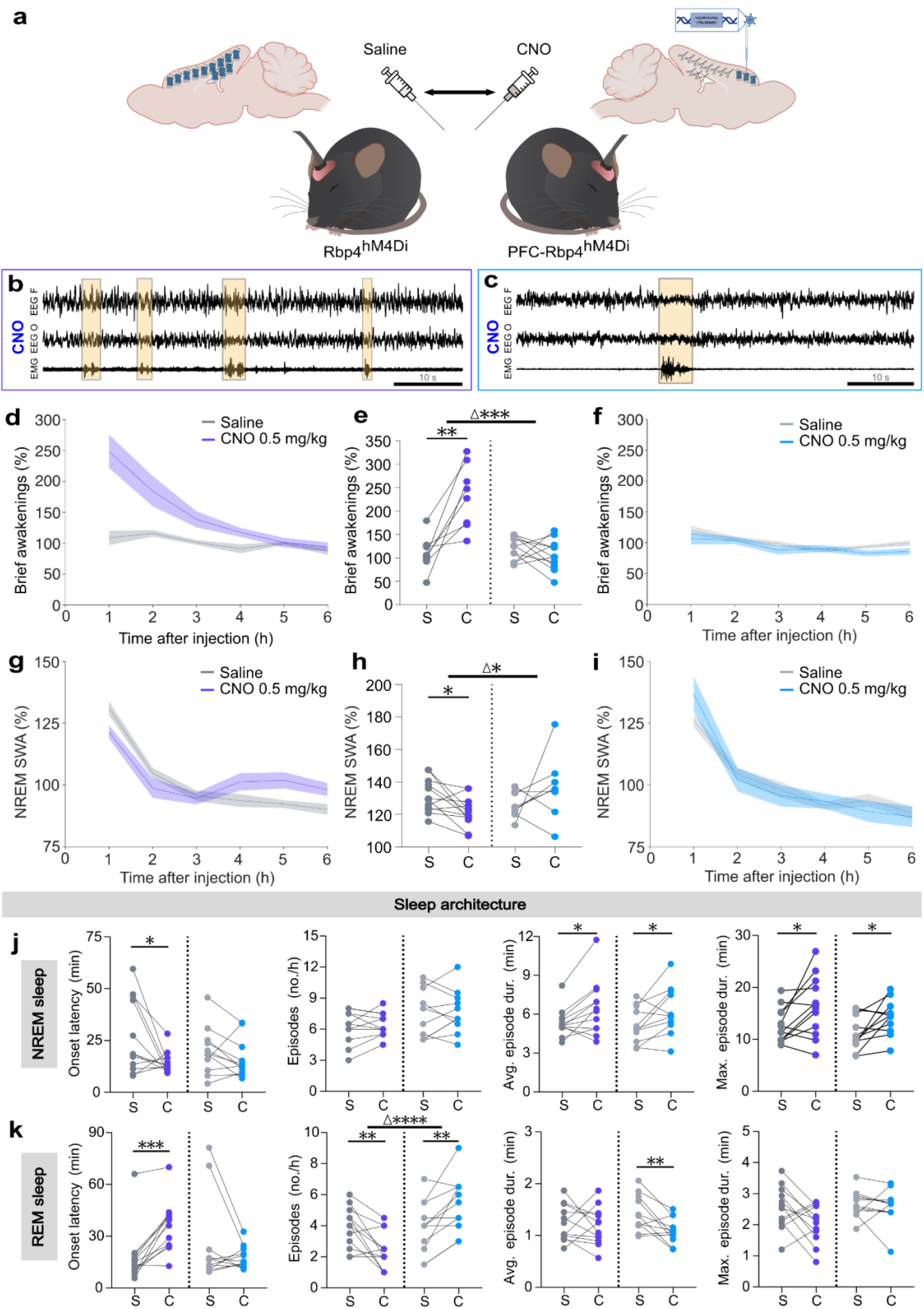
Global but not local cortical inhibition of *Rbp4-Cre+* neurons disrupts NREM sleep and reduces slow-wave activity. **(a)** Schematic of the comparison of electrophysiological recordings in Rbp4^hM4Di^ (‘global’, left) and PFC-Rbp4^hM4Di^ (local PFC, right) mice, following counterbalanced intraperitoneal (i.p.) injections of saline and 0.5 mg/kg clozapine-*N*-oxide (CNO) at light onset (ZT=0). **(b and c)** Representative electroencephalogram (EEG, Frontal F and Occipital O) and electromyogram (EMG) traces after CNO injections in **(b)** global Rbp4^hM4Di^ versus **(c)** local PFC-Rbp4^hM4Di^ mice, with brief awakening examples highlighted in yellow. **(d-f)** Time courses of brief awakenings (BA; <16 s movement during sleep; number per hour total sleep time, expressed as percentage of saline recording average) during the first 6 hr after saline (S) and CNO (C) in **(d)** global Rbp4^hM4Di^ versus **(f)** local PFC-Rbp4^hM4Di^ mice, with **(e)** the corresponding number of BA’s in the first hour post-i.p. Injection. **(g-i)** Time courses of EEG NREM slow-wave activity (SWA, % saline recording average) for the first 6 hr after saline (S) and CNO (C) in **(g)** global Rbp4^hM4Di^ versus **(i)** local PFC-Rbp4^hM4Di^ mice, with **(h)** the corresponding % of NREM SWA in the first hour post-i.p. injection. **(j and k)** Sleep architecture during **(j)** NREM and **(k)** REM sleep in hours 1+2 post-injection of saline (S) and CNO (C) in global Rbp4^hM4Di^ (purple) versus local PFC-Rbp4^hM4Di^ mice (cyan). Note that all time analyses were aligned from 15 min post-injection (as T0) to ensure full DREADD activation and are visualised as mean ± SEM (shaded). Asterisks indicate comparisons with significant differences (*p<0.05; **p<0.01; ***p<0.001, ****p<0.0001) for analyses with significant main effects. n=9 Rbp4^hM4Di^ for panels b and d, n=11 PFC-Rbp4^hM4Di^ for panels c and f, n=12 Rbp4^hM4Di^ for panel g, n=8 PFC-Rbp4^hM4Di^ for panel i, n=12 Rbp4^hM4Di^ and n=11 PFC-Rbp4^hM4Di^ for panels e, h, j and k. C: CNO; CNO: Clozapine-*N*-oxide; DREADD: Designer Receptors Exclusively Activated by Designer Drugs; EEG: Electroencephalogram; EMG: Electromyogram; i.p.: intraperitoneal; NREM: Non-rapid eye movement sleep; PFC: Prefrontal cortex; REM: Rapid eye movement sleep; S: Saline; SWA: Slow-wave activity; TST: Total sleep time; T0: timepoint of analysis onset 15 min post-injection; ZT: Zeitgeber time.

We first analysed differences in sleep fragmentation, as measured by the number of brief awakenings (short bouts <16 s of wake-like EEG/EMG activity during sleep). In global Rbp4^hM4Di^ mice, the number of brief awakenings (expressed as % of 6 hr saline recording average) was substantially increased in the CNO condition compared to saline (saline: 108.7±34.98%, CNO 0.5 mg/kg: 248.8±81.37%; *t*(8) = 4.72, *p* = 0.002) (Fig. 3b,d,e) and this increase lasted approximately 3-4 hours. In contrast, the frequency of brief awakenings was unaltered during the DREADD manipulation in PFC-Rbp4^hM4Di^ mice compared to saline (saline: 119.9±24.71%, CNO 0.5 mg/kg: 107.0±32.86%; *t*(10) = 0.97, *p* = 0.36) (Fig. 3c,e,f). When directly contrasted, there was a significant difference in the change in brief awakenings between the Rbp4^hM4Di^ and PFC-Rbp4^hM4Di^ groups (Δ_global inhibitory_: +140.0±89.02%, Δ_local inhibitory_: −12.93±44.25%; *t*(11.19) = 4.701, *p* = 0.001) (Fig. 3e).

We next analysed NREM SWA (0.5-4 Hz; expressed as % of 6 hr saline recording average), an established marker of sleep intensity^1,3,18^. Despite greater NREM sleep time in the global Rbp4^hM4Di^ group following CNO (Fig. 1g,h), we observed an altered NREM SWA profile in these mice; that is, SWA was significantly reduced in the first hour post-DREADD activation (saline: 131.5±10.38%, CNO 0.5 mg/kg: 121.6±9.20%; *t*(11) = 2.883, *p* = 0.015) (Fig. 3h). In contrast, SWA in PFC-Rbp4^hM4Di^ mice during DREADD-induced NREM sleep did not differ from the saline condition (saline: 126.10±8.06%, CNO 0.5 mg/kg: 136.80±19.8%; *t*(7) = 1.244, *p* = 0.254) (Fig. 3h,i), indicating that sleep intensity was maintained. When NREM SWA was directly contrasted between the global Rbp4^hM4Di^ and PFC-Rbp4^hM4Di^ groups, the changes in SWA between CNO and saline were significantly different (Mdn_Δglobal inhibitory_ = −6.72%, IQR_Δglobal inhibitory_ = 18.43%, Mdn_Δlocal excitatory_ = +11.49%, IQR_Δlocal excitatory_ = 26.47%; *U* = 16, *p* = 0.012) (Fig. 3h). Interestingly, NREM SWA appeared to rebound in the global Rbp4^hM4Di^ group approximately 4-6 hours post-DREADD activation (Fig. 3g), typical of a response to sleep deprivation^12,18^. Analysis of the entire 0.5-30 Hz spectral range additionally revealed a DREADD-induced suppression of higher frequencies (>3.5 Hz) during NREM sleep and left-shifted REM sleep theta peak in global Rbp4^hM4Di^ mice (Suppl. Fig. 3a). PFC-Rbp4^hM4Di^ mice had a mild reduction of frequencies above 5-6 Hz during NREM sleep and unaltered REM sleep spectra post-CNO compared to saline (Suppl. Fig. 3c). No differences in NREM SWA nor any other NREM/REM sleep spectral features were observed between saline and CNO in the DREADD-free R26^LSL-hM4Di^ control group (Suppl. Fig. 3b).

To further investigate globally- versus locally-induced sleep, we analysed NREM and REM architecture differences. All global Rbp4^hM4Di^ mice fell asleep within 30 minutes of CNO injections and had shorter NREM latencies (saline: 26.76±17.85 min, CNO 0.5 mg/kg: 14.21±5.24 min; *t*(11) = 2.414, *p* = 0.034) and, compared to saline, had longer NREM episodes (Mdn_saline_ = 5.09 min, IQR_saline_ = 0.7 min, Mdn_CNO_ = 6.33 min, IQR_CNO_ = 2.78 min; *W* = 58, *p* = 0.021) (Fig. 3j). REM onset was significantly delayed (Mdn_saline_ = 14.37 min, IQR_saline_ = 9.32 min, Mdn_CNO_ = 37.73 min, IQR_CNO_ = 17.83 min; *W* = 76, *p* = 0.001) and fewer REM episodes occurred per hour (saline: 3.88±1.32/hour, CNO 0.5 mg/kg: 2.42±1.2/hour; *t*(11) = 3.554, *p* = 0.005) (Fig. 3k). This is consistent with the observed NREM-promoting and REM-suppressing effects observed in the analysis of sleep amount (Fig. 1f,g). In contrast, latencies to NREM and REM sleep did not change in the PFC-Rbp4^hM4Di^ mice between saline and CNO, but the length of NREM sleep episodes (saline: 5.18±1.39 min, CNO 0.5 mg/kg: 6.27±1.87 min) (Fig. 3j) and number of REM episodes (saline: 3.91±1.72/hour TST, CNO 0.5 mg/kg: 5.36±1.79/hour TST) were increased (*p* = 0.034 and *p* = 0.006, respectively), and the average length of REM episodes was reduced (saline: 1.43±0.35 min, CNO 0.5 mg/kg: 1.09±0.25 min; *t*(10) = 3.263 *p* = 0.009) (Fig. 3k). Sleep architecture was unaffected in PFC-Rbp4^hM3Dq^ mice (Suppl. Fig. 2).

Overall, acute inhibition of cortical *Rbp4-Cre+* neurons on both global and local spatial scales increased NREM sleep amount, but globally-induced sleep was less consolidated with reduced intensity and strong architectural differences, whereas PFC-induced sleep left key physiological sleep features largely intact.

### ‘Local’ occipital chemogenetic inhibition and excitation of *Rbp4-Cre+* neurons bidirectionally alters sleep architecture but not overall sleep/wake amounts

To determine whether local *Rbp4-Cre+* neuronal activity in cortical regions beyond the PFC similarly altered sleep, we next applied the same viral-vector strategy for chemogenetic inhibition/excitation to *Rbp4-Cre+* neurons in the OCC (Fig. 4a,b). This was informed by two recent studies demonstrating a role for layer 2/3 pyramidal neurons in the retrosplenial and primary visual cortices in the initiation and maintenance of REM sleep^22,26^.

**Figure 4:**
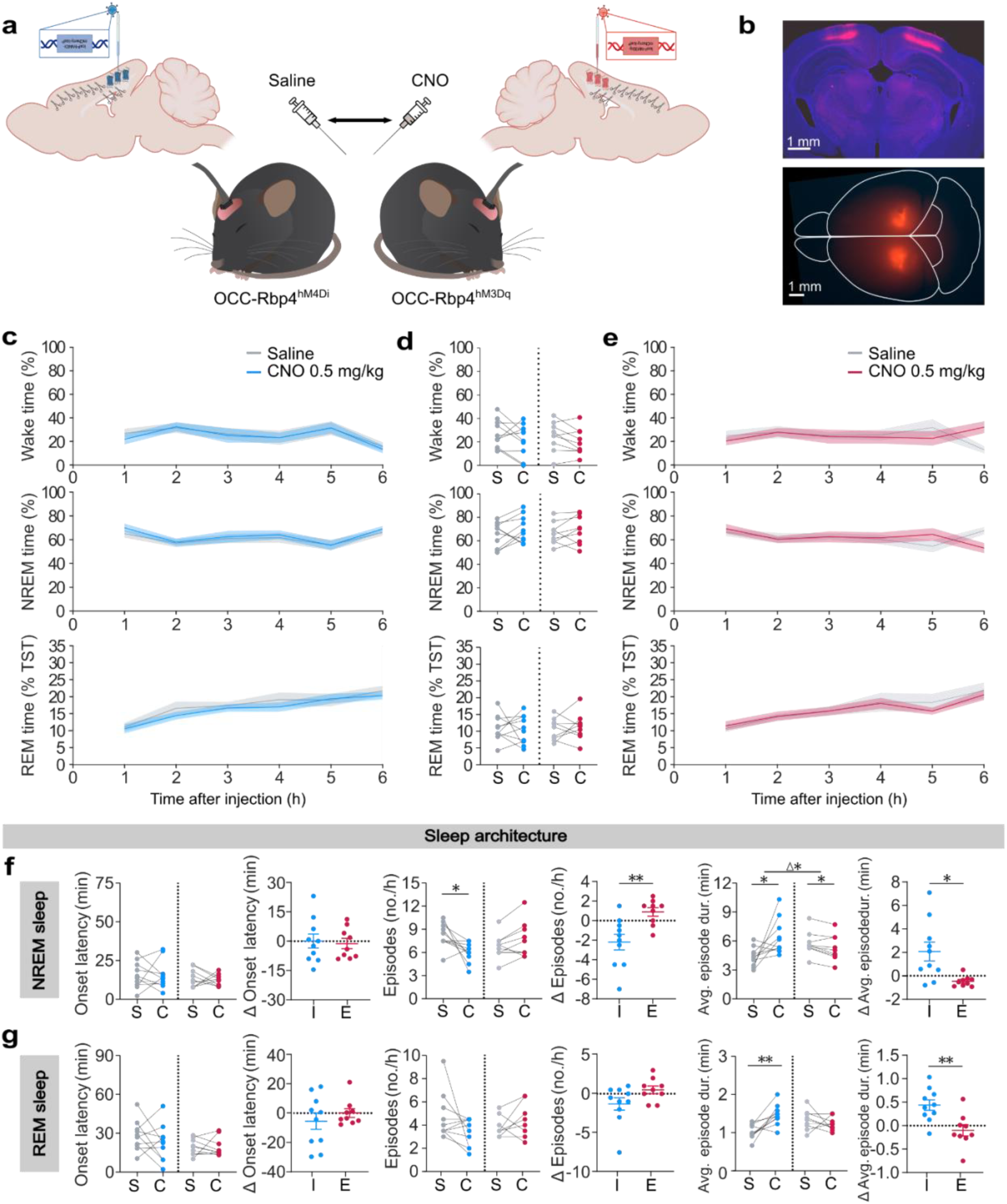
Chemogenetic inhibition/excitation of occipital cortical layer 5 pyramidal neurons bidirectionally modifies sleep architecture, without impacting sleep amount. **(a)** Schematic of viral vector injections (left: inhibitory hM4Di; right: excitatory hM3Dq) for localised DREADD expression in the OCC of Rbp4^Cre^ mice. **(b)** Coronal brain section (top) and dissection microscope image (bottom) showing OCC DREADD expression. Scale bars, 1000 µm. **(c and e)** Time courses of Wake, NREM, and REM sleep in the first 6 hr post-intraperitoneal (i.p.) injection of saline and CNO in **(c)** OCC-Rbp4^hM4Di^ and **(e)** OCC-Rbp4^hM3Dq^ mice, with **(d)** the corresponding percentages of time spent in each vigilance state during the first hour post-i.p. injection at light onset (ZT=0). Sleep architecture during NREM **(f)** and REM **(g)** sleep in hours 1+2 post-injection of saline and CNO in OCC-Rbp4^hM4Di^ versus OCC-Rbp4^hM3Dq^ mice, with direct comparisons of the difference (Δ) between CNO and saline conditions in the two groups (I: Inhibitory; E: Excitatory). Rapid eye movement (REM) sleep is presented as a percentage of total sleep time (% TST). Note that all time analyses were aligned from 15 min post-injection (as T0) to ensure full DREADD activation and are visualised as mean ± SEM (shaded). Asterisks indicate post-hoc comparisons with significant differences (*p<0.05; **p<0.01) for analyses with significant main effects. n = 10 OCC-Rbp4^hM4Di^ for panel c, n=9 OCC-Rbp4^hM3Dq^ for panel e, n = 10 OCC-Rbp4^hM4Di^ and n=9 OCC-Rbp4^hM3Dq^ for panels d, e and f. C: CNO, Clozapine-N-oxide; DREADD: Designer Receptors Exclusively Activated by Designer Drugs; E: Excitatory DREADD; I: Inhibitory DREADD; i.p.: intraperitoneal; NREM: Non-rapid eye movement sleep; OCC: Occipital cortex; REM: Rapid eye movement sleep; S: Saline; TST: Total sleep time; T0: timepoint of analysis onset 15 min post-injection; ZT: Zeitgeber time.

In OCC-Rbp4^hM4Di^ and OCC-Rbp4^hM3Dq^ mice, time spent in Wake, NREM and REM sleep did not change following DREADD activation compared to saline (*p* > 0.1 in all cases) (Fig. 4c-e), nor did latencies to NREM or REM sleep (*p* > 0.1 for all comparisons) (Fig. 4f,g), nor did NREM SWA or any other spectral frequencies during NREM or REM sleep (Suppl. Fig. 3e,f). Interestingly, however, OCC-Rbp4^hM4Di^ mice exhibited a significant reduction in the number – but increase in the duration – of NREM episodes (NREM episode number – saline: 8.25±1.6/hour, CNO 0.5 mg/kg: 6.05±1.28/hour; *t*(9) = 2.762, *p* = 0.022; NREM episode duration – saline: 4.35±1 min, CNO 0.5 mg/kg: 6.42±1.85 min; *t*(9) = 2.587, *p* = 0.029). An inverse tendency was observed following excitation (NREM episode number – saline: 6.83±1.6/hour, CNO 0.5 mg/kg: 7.72±2.33/hour; *t*(9) = 2.762, *p* = 0.0732; NREM episode duration – saline: 5.75±1.24min, CNO 0.5 mg/kg: 5.29±1.34 min; *t*(8) = 3.116, *p* = 0.014) (Fig. 4f). The number of REM episodes remained unchanged in both groups (p > 0.1 in both cases) but average REM episode duration was significantly increased following CNO administration in OCC-Rbp4^hM4Di^ mice (saline: 1.04±0.19 min, CNO 0.5 mg/kg: 1.48±0.27 min; *t*(9) = 3.952, *p* = 0.003) and decreased in OCC-Rbp4^hM3Dq^ mice (saline: 1.34±0.26 min, CNO 0.5 mg/kg: 1.24±0.18 min; *t*(8) = 0.847, *p* = 0.42) (Fig. 4g). Direct comparison of the difference between CNO and saline conditions in the two groups demonstrated a bidirectional change for NREM episode number (Δ_inhibitory_: −2.2±2.52/hour; Δ_excitatory_: +0.89±1.29/hour; *t*(13.72) = 3.41, *p* = 0.004) (Fig. 4f,), episode duration (Δ_inhibitory_: +2.07±2.53min; Δ_excitatory_: −0.46±0.45min; *t*(9.62) = 3.113, *p* = 0.012), as well as REM episode duration (Δ_inhibitory_: +0.37±0.3min; Δ_excitatory_: - 0.1±0.36min; *t*(15.51) = 3.05, *p* = 0.008) (Fig. 4g).

## Discussion

The cerebral cortex exerts a strong influence over many complex behaviours – from emotion regulation to motor planning to social interactions. From this perspective, the emerging notion that the cerebral cortex also regulates key sleep features – including sleep preparation and initiation^21^, NREM and REM sleep amounts^10,22,23,25–28^, and synaptic strength-dependent sleep pressure^24^ – is unsurprising. Here we begin to resolve the contributions of different spatial scales of cortical activity upon sleep, by investigating the effects of global versus local manipulations of a distinct cortical neuronal population.

First, we have shown that cortex-wide, global chemogenetic inhibition of *Rbp4-Cre+* neurons can strongly induce sleep. However, this sleep had an unusually high number of brief awakenings and suppression of SWA sleep, indicating increased sleep fragmentation and reduced sleep intensity, respectively. REM sleep was also strongly suppressed. We then observed that targeted, ‘local’ chemogenetic inhibition and excitation of these neurons in the PFC bidirectionally altered NREM sleep amount to a similar extent as in the global group – despite implicating a substantially smaller number of cortical neurons – and left REM sleep, brief awakenings, and sleep spectra largely in-tact. Interestingly, local chemogenetic manipulations of the same cell type in the OCC left overall sleep/wake amounts unaffected but bidirectionally altered several sleep architecture parameters.

There are several possible explanations for the strong sleep-promoting effects of inhibiting *Rbp4-Cre+* neurons in the cerebral cortex, and specifically in the PFC. First, PFC L5 pyramidal cells could have a top-down influence over sleep centres in the hypothalamus and brainstem via projections similar to those of PFC*^Sst^*^-GABA^ neurons, which trigger sleep preparatory behaviours and induce sleep through projections to the lateral preoptic area and lateral hypothalamus^21^. Second, L5 pyramidal neurons could initiate slow oscillations that propagate through the neocortex^29^ and synchronise global cortical activity such as in general anaesthesia^30^; acute chemogenetic inhibition/excitation might facilitate/suppress such activity patterns in cortical networks, and subsequent changes in sleep/wake state. Finally, intra-cortically projecting layer 5 pyramidal neurons may engage in crosstalk with PFC*^Sst^*^-GABA^ neurons or other cortical sleep-regulatory cells, such as the transcriptomically re-defined population of *SST/Chodl/Nos1/Tacr1* (SCNT) neurons^31^, which promote cortical synchrony and sleep^28^. Interestingly, in parallel to our work, a recently discovered PFC layer 5 pyramidal neuronal subtype, *Colgalt2* neurons, are being investigated for their NREM sleep- and anaesthesia-promoting (NAP) properties^32^.

Beyond providing evidence for the hypothesis that ‘global’ sleep – that is, sleep as a brain state and behaviour – can passively emerge from ‘local’ sleep (i.e., localised sleep-like activity in the cortex^11,33^), our data suggests that the global state can be actively determined by changes in local activity. Importantly, the aspects of sleep altered by local activity changes appear highly dependent on the cortical region and spatial extent of the manipulation. For example, we observed that the sleep induced by our cortex-wide inhibition had unphysiological features – suppressed REM, highly frequent brief awakenings, reduced NREM SWA –, while the sleep induced by local chemogenetic inhibition of only a small subpopulation in the PFC left naturalistic sleep features intact. The OCC manipulations influenced sleep differently again – they altered sleep architecture but not sleep/wake amounts – indicating that different areas of cortex may be feeding different information into sleep regulation. Together, our data suggest that global cortical signals implicating neurons with diverse inputs and outputs, may be producing complex and possibly conflicting interactions that result in abnormally regulated sleep, while local activity changes in sleep-regulatory populations – in the right location – may afford finer control.

Notably, classical hypnotics like benzodiazepine and ‘Z-drugs’ (benzodiazepine-receptor agonists) act globally to increase sleep time but disrupt sleep architecture and electrophysiology and impair core functions of sleep^34,35^. This may represent a shared disadvantage of systemic sleep-promoting approaches that alter cortex- or even brain-wide activity, like in our global Rbp4^hM4Di^ mice. Many pharmacological approaches to induce sleep which operate this way come at the cost of detrimental health consequences^36^, highlighting how increasing the amount of sleep alone may be insufficient to harvest sleep’s benefits. More targeted manipulations of sleep-regulatory areas in humans may make it possible to induce sleep with preserved physiological features, such as in our local PFC manipulation in mice. Clinically, neocortical activity and related networks can now be effectively modulated with novel non-invasive brain stimulation approaches that afford high spatial precision. Subregions of the PFC are already evidence-based targets for neuromodulation in depression^37^ and can now be precisely stimulated with individualised parameters to enhance SWA during NREM sleep^38^. Our data supports the PFC as an accessible, local neocortical target with the capacity to promote naturalistic sleep in a way that has not yet been achieved through global interventions.

## Methods

### Mice

The following types of mice were used: *Rbp4-Cre+* mice *(Tg(Rbp4-Cre)KL100Gsat/Mmucd)* expressing Cre-recombinase in layer 5 pyramidal neurons of the neocortex and dentate gyrus granule cells of the archicortex^39,40^, kindly provided by R.M. Bruno (University of Oxford, UK); R26^LSL-hM4Di^ mice (*B6.129-Gt(ROSA)26Sortm1(CAG-CHRM4*,-mCitrine)Ute/J*, Jackson Laboratory stock #026219) for Cre recombinase-inducible expression of the HA-tagged DREADD receptors hM4Di and the yellow fluorescent protein mCitrine; and a cross of the first two strains (abbreviated here as Rbp4^hM4Di^ mice). The *Rbp4-Cre+* mouse line was bred to homozygosity and congenic on the C57BL/6 background, and the R26^LSL-hM4Di^ mouse line was supplied by the Jackson Laboratory on a C57BL/6 background and maintained homozygously in our local colony. A total of 73 young adult male mice were used: 14 Rbp4^hM4Di^; 9 R26^LSL-hM4Di^; and 50 Rbp4^Cre^ (13 PFC-Rbp4^hM4Di^, 10 PFC-Rbp4^hM3Dq^, 14 OCC-Rbp4^hM4Di^, and 13 OCC-Rbp4^hM3Dq^). All mice were housed on a 12 h light/12 h dark cycle, at constant temperature and humidity and with *ad libitum* access to food and water.

### Stereotaxic surgery

#### EEG/EMG implants

All surgeries were aseptic and performed under isoflurane anesthesia as previously described^34^. Two electroencephalogram (EEG) screw electrodes were placed at mediolateral (ML) +2.0 mm/anteroposterior (AP) +2.0 mm (frontal derivation) and ML +2.5/AP −3.5 mm (occipital derivation), relative to bregma. Where multiple viral injections were done (see below), the position of the EEG screw near the injection sites was slightly adjusted (frontal screw coordinate to ML +2.2 /AP +1.8 mm; occipital screw coordinate to ML +2.5/AP −4 mm). A reference screw electrode was placed over the cerebellum at approximately ML 0 /AP −1.5 mm. Two electromyography (EMG) wire electrodes were inserted bilaterally into the neck extensor muscles.

#### Viral injections

Local expression of inhibitory and excitatory DREADD receptors in the prefrontal (PFC) or occipital cortex (OCC) was achieved by injecting the viruses *pAAV8-hSyn-DIO-hM4D(Gi)-mCherry* (Addgene plasmid #44362) and *pAAV8-hSyn-DIO-hM3D(Gq)-mCherry* (Addgene plasmid #44361), respectively, in Rbp4^Cre^ mice. 200 nL of virus was bilaterally injected at 1-4 sites per hemisphere (PFC: ML ±0.65 /AP +2.6 mm; ML ±1.5 /AP +2.6 mm; ML ±0.65 /AP +1.9 mm; ML ±1.5/AP + 1.9 mm; OCC: ML ± 1.7 /AP - 3.1 mm; ML ± 2.6/AP - 3.1 mm; ML ± 1.9 /AP - 2.4 mm; all relative to bregma), across two depths (dorsoventral (DV), PFC: −1.5 and −0.8 mm; OCC: −0.5 and −0.8 mm, relative to the brain surface). Medial injections in the PFC (that is, injections done at ML ± 0.65 mm) were angled six degrees inwards to avoid laceration of the superior sagittal sinus and related veins. The virus infusion rate was 40 nL/min using a Nanoject II (Drummond Scientific). There was at least 3 weeks’ wait between viral injections and the first recordings to ensure adequate viral transgene expression.

### Chronic electrophysiological recordings

Mice were acclimatised to custom-built Plexiglas recording chambers (20.3 x 32 x 35 cm) and tethered recording conditions for at least 3 days prior to the start of experiments. The chambers were placed in sound-attenuated and light-controlled Faraday cages (Campden Instruments Ltd., London, UK). EEG/EMG signals were recorded with the 128-channel Neurophysiology Recording System (Tucker-Davis Technologies Inc., Alachua, FL, USA) and the electrophysiological recording software Synapse. Frontal and occipital EEG derivations were referenced to a screw electrode over the cerebellum. Raw electrophysiological signals were filtered (0.1-100 Hz) and amplified with a PZ5 NeuroDigitizer (Tucker-Davis Technologies Inc., Alachua, FL, USA), and stored at a sampling rate of 305 Hz on a local computer.

### Chemogenetic manipulations

To chemogenetically modulate cortical neurons, mice were intraperitoneally injected with clozapine*-*N-oxide dihydrochloride (CNO) and saline (control). The CNO solution used was prepared from clozapine-N-oxide dihydrochloride powder (Tocris, Bio-Techne LTD, Abingdon, UK, catalog no.: 6329) with sterile 0.9% saline and passed through a Millipore filter. The main experiments comprised counterbalanced conditions of 0.5 mg/kg CNO and saline at light onset (zeitgeber time 0), when homeostatic sleep pressure is typically at a moderate level. To assess a potential dose-dependency and to determine optimal dosage, the Rbp4^hM4Di^ mice received up to four different conditions at light onset (saline: n = 14; 0.5 mg/kg CNO: n = 12; 1 mg/kg CNO: n = 10; 5 mg/kg CNO: n = 9). In three OCC-Rbp4^Cre^ mice, the initial 0.5 mg/kg CNO vs. saline injections were repeated due to technical issues and data from the repeat conditions were used.

#### CNO considerations

While generally considered biologically inert, CNO undergoes partial back-conversion to clozapine^42^ and has off-target binding at endogenous neurotransmitter receptors^43^. We previously systematically tested the effects of different CNO doses (1, 5 and 10 mg/kg) on sleep in wild-type (C57BL/6) mice that do not express hM3Dq or hM4Di DREADD receptors^41^. We reported that CNO elicits a dose-dependent effect on several sleep parameters at medium and high doses (5 and 10 mg/kg) of this water-soluble CNO product. More specifically, it causes a mild REM suppression, alters individual NREM and REM bout number and duration, and increases sleep state stability and sleep continuity^41^. For low-dose CNO (1 mg/kg) we found no significant effects on sleep amount, sleep electrophysiology, and nearly all other assessed sleep architectural parameters reported in this current study, but observed some indications of small effects on NREM and REM latencies and brief awakenings. Here, we used an even lower dose of CNO (0.5 mg/kg), in line with other recent DREADD-based sleep studies in mice which show no DREADD-independent effects in their control groups, but a sufficient activation of DREADDs to test sleep-wake regulation^44^. Here we additionally provide data from a DREADD-free control group (Suppl. Fig. 1, 3b) injected with the same CNO dose and product to exclude off-target CNO effects in our findings.

### Histology

Mice were deeply anaesthetised with pentobarbital and transcardially perfused with 0.01 M PBS followed by 4% formaldehyde made up from 37% stock solution (product code F8775, Sigma-Aldrich). Brains were removed and fixed in PFA overnight then stored in PBS. Whole brains were imaged on a fluorescence dissection microscope equipped with a Leica DFC490 digital camera to assess the regional spread of viral expression before being sliced for histological assessment into 50 µm coronal sections using a Leica VT1000S vibrating microtome. Sections were stained with DAPI before being mounted onto glass slides and coverslipped. Fluorescence images were taken with a Leica DMR upright fluorescence microscope equipped with a Leica DFC500 digital camera. Images were analyzed and merged and scale bars were added using FIJI ImageJ (v2.9.0). All figures were created using GraphPad Prism (v11.0.1) and then styled in Inkscape (v1.0.2, Inkscape Project 2020; https://inkscape.org). The results across all animals were similar to those presented in the representative images. Based on a predetermined exclusion criteria of unilateral or no viral expression, one PFC-Rbp4^hM4Di^, four OCC-Rbp4^hM4Di^ and four OCC-Rbp4^hM3Dq^ mice were excluded from all analyses. One PFC-Rbp4^hM4Di^ mouse was additionally excluded due to a cortical structural anomaly.

### Electrophysiological data processing and analysis

Filtered and amplified EEG/EMG signals were resampled at 256 Hz offline using custom MATLAB (The MathWorks Inc., Natick, MA, USA, v. R2023b) scripts and converted into the European Data Format (EDF) via the open-source software Neurotraces. All files were sleep scored in a blinded, semi-automated fashion with SleepSign for Animals (v3.3.6.1602, SleepSign Kissei Comtec). EEG/EMG recordings were partitioned into 4-second epochs and automatically pre-annotated with a vigilance state assignment (Wake, NREM, or REM). Annotations were manually checked to ensure mainly that artefacts, movements, and vigilance state transitions were correctly scored, based on visual inspection of the frontal and occipital EEG derivations and EMG traces.

Epochs with recording artifacts arising from gross movements, chewing or external electrostatic noise were still assigned to the respective vigilance state but not included in the spectral analysis. EEG power spectra were computed using a fast Fourier transform routine (Hanning window) with a 0.25-Hz resolution and exported in the frequency range between 0 and 30 Hz for spectral analysis. NREM SWA (%) was calculated as a percentage of the average NREM SWA (the spectral power in the frequency bins between 0.5 and 4 Hz) relative to the 6-hour observation time window following saline injections. Brief awakenings (BAs, %) were calculated as a percentage of the number of BAs, per hour of total sleep time, relative to the 6-hour observation time window following saline injections. All other NREM/REM architecture analyses were based on EEG/EMG recordings scored according to criteria previously detailed^41^. Six animals included in the vigilance state analysis had to be excluded from the analysis of brief awakenings due to insufficient EMG quality (n=3 Rbp4^hM4Di^, n=1 R26^LSL-hm4Di^, n=2 PFC-Rbp4^hM3Dq^). Due to reduced EEG quality, six animals included in vigilance state analysis were excluded from frontal SWA analysis (n=1 R26^LSL-hM4Di^, n=3 PFC-Rbp4^hM4Di^, n=2 OCC-Rbp4^hM3Dq^).

### Statistics

Data were analysed using MATLAB (versions R2023b and 2025b; The MathWorks Inc, Natick, MA, USA) and GraphPad Prism (version 11.0.1 for Windows; GraphPad Software, San Diego, CA, USA, https://www.graphpad.com/). Data are presented as mean ± SD and are rounded to two decimal points, unless stated otherwise. Statistical significance was set at p < 0.05 and all tests were two-sided. Except where incorporated into post-hoc tests, reported p-values were not adjusted for multiple comparisons. Reported p-values and test statistics are rounded to three decimals.

To accommodate occasional missing repeated measurements, differences between saline and varying CNO doses were assessed separately for each vigilance state using mixed-effect models (restricted maximum likelihood method) with Geisser-Greenhouse correction. Treatment conditions were specified as repeated measures within subjects. Dunnett’s post-hoc test was applied where appropriate, with post-hoc tests being restricted to reference group comparison against saline. Model residuals were evaluated for normality using Shapiro-Wilk tests in combination with visual inspection of Q–Q and residual plots.

Comparisons between saline and 0.5 mg/kg CNO conditions within the global Rbp4^hM4Di^, PFC-Rbp4^hM4Di^, PFC-Rbp4^hM3Dq^, OCC-Rbp4^hM4Di^ and OCC-Rbp4^hM3Dq^ groups were performed using paired t-tests or Wilcoxon signed-rank tests when parametric assumptions were not met. In such cases, median and interquartile range was reported instead of mean ± SD. Between-group differences (calculating Δ as the within-subject difference between CNO and saline conditions) were assessed via unpaired Welch’s t-tests or Mann-Whitney U tests where appropriate. These between-group comparisons were, on all occasions, preceded by control comparisons of the two saline conditions via unpaired Welch’s t-tests or Mann-Whitney tests, when appropriate. Comparability of saline conditions can be considered maintained in the analyses unless explicitly stated otherwise. For nonparametric, rank-based tests, ties were taken into account throughout all analyses, as per software default settings.

Note that all time window details above were defined relative to 15 min after intraperitoneal (i.p.) injection, informed by previous pharmacokinetic work showing that this is the time point at which CNO reaches peak plasma, brain and CSF levels^43^, and by examples from similar chemogenetic studies which allow up to 20 min for the onset of DREADD-mediated effects^44^.

### Ethical approval

All experiments were performed in accordance with the UK Home Office Animal Scientific Procedures Act (1986) under personal and project licenses granted by the United Kingdom Home Office. Ethical approval was provided by the Local Ethical Review Panel at the University of Oxford. Animal holding and experimentation was located at the Biomedical Sciences Building, University of Oxford.

## Author contributions

L.B.K.: conceptualisation, funding, electrophysiology, histology, sleep scoring, data analysis, supervision, manuscript writing and editing. J.D.H.: conceptualisation, electrophysiology, histology, sleep scoring, data analysis, manuscript writing and editing. A.H.S.: histology, supervision, manuscript writing and editing. C.M.: electrophysiology, histology, sleep scoring, manuscript writing and editing, data analysis. L.M.: electrophysiology, histology, sleep scoring, manuscript editing, data analysis. A.B.S.: data analysis, manuscript writing and editing. C.J.A.: conceptualisation, supervision, manuscript writing and editing. Z.M.: conceptualisation, funding, supervision, manuscript editing. V.V.V.: conceptualisation, funding, supervision, manuscript writing and editing.

## Acknowledgements

We thank the members of the Vyazovskiy, Molnár and Miesenböck labs for discussions and feedback on the project. In particular we thank Dr. Linus Milinski for detailed feedback on the figures and text. We thank Liliana Mendes da Paz, James Metcalf, and Laura Thomas of the Oxford Mouse Sleep Core (OMSC) for their technical support, and the Oxford Biomedical Sciences Level 1 and 2 teams for taking care of the animals. We thank Madison Bartley and Prof. Randy Bruno for the provision of breeder pairs for the Rbp4^Cre^ mouse line. We thank Louise Aarons for assistance with surgeries, electrophysiology experiments, and sleep scoring as part of a student project. We thank Silav Sadoun for supporting histology and data processing. We thank Dr. Natalie Hauglund for the mouse illustration and Dr. Clifford Talbot for helping with statistical analyses. Finally, we thank L.B.K.’s Sir Henry Wellcome Fellowship sponsors Prof. Chiara Cirelli, Prof. Antoine Adamantidis, and Prof. Gero Miesenböck for their ongoing advice and support.

We thank the following funders for their financial support: L.B.K. was supported by a Wellcome Trust Doctoral Studentship in Neuroscience (203971/Z/16/Z) and is currently supported by a Wellcome Trust Sir Henry Wellcome Fellowship (224083/Z/21/Z), and the Exeter College Staines Medical Research Fellowship. J.D.H. is supported by an Australian Ramsay Postgraduate Scholarship, James Fairfax Oxford-Australia Scholarship, and previously the Oxford Clarendon Fund. C.M. is supported by the German Academic Scholarship Foundation, BMEP Stipend (University Clinic Jena), and an Oxford Berlin Research Partnership (Early Career Researcher Mobility). L.M. is supported by the German Academic Scholarship Foundation. C.J.A. was supported by MRC project MR/Z504567/1 and MRC project MR/S01134X/1. Z.M. and V.V.V. are supported by a BBSRC grant (BB/X008711/1). Z.M. is further supported by an MRC Project Grant (G00900901) and a Research Grant from St John’s College Research Centre (21138077). V.V.V. is further supported by a Wellcome Trust grant (227093/Z/23/Z). No artificial intelligence was used for data analysis or writing of this manuscript. The large language model ChatGPT (version 5, OpenAI, 2025) was used to simplify and annotate Matlab scripts previously written by L.B.K and V.V.V. and to adapt the colour scheme of illustrations.

None of the authors declare a conflict of interest.

## Supplementary Materials

**Suppl. Fig. 1:**
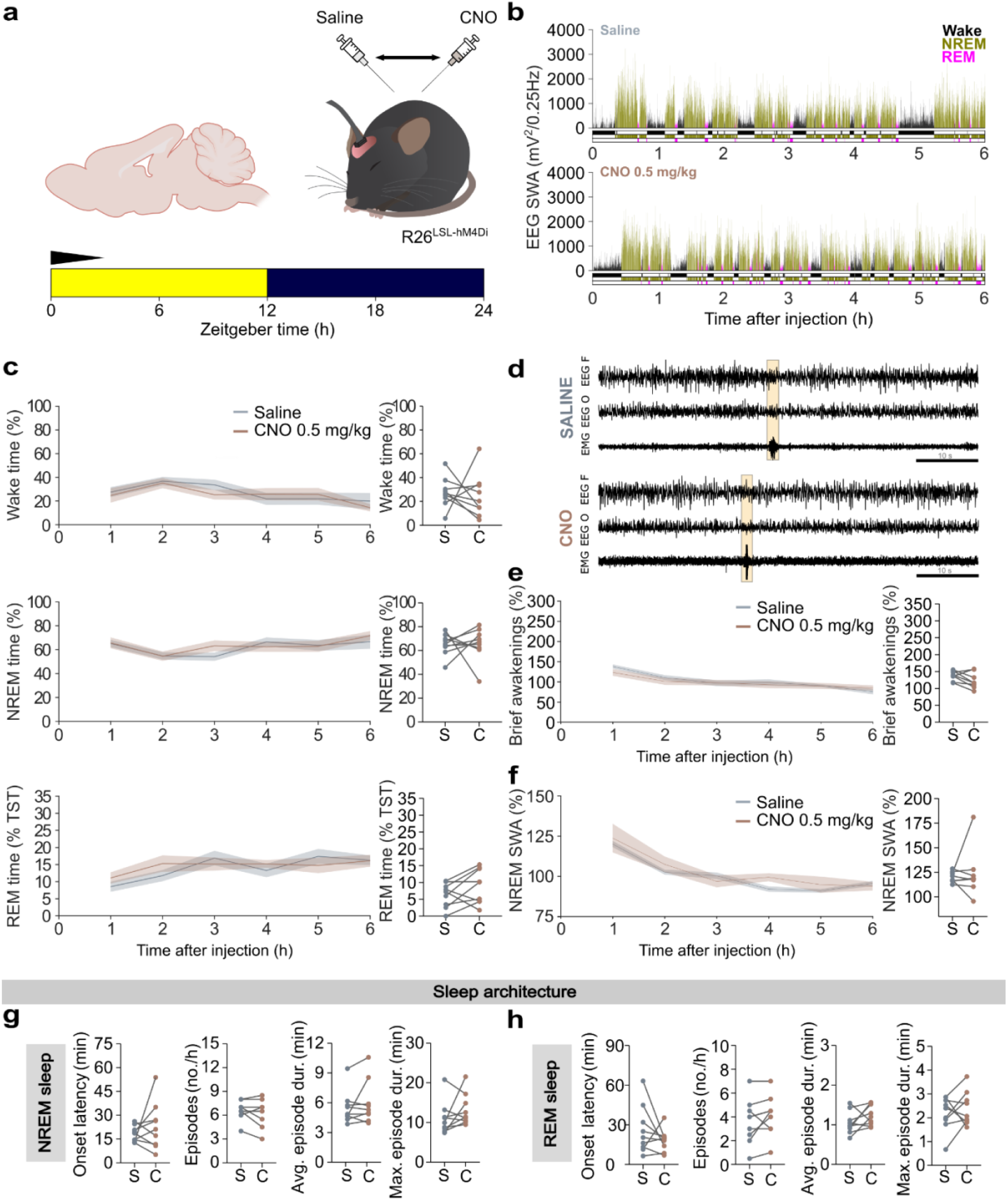
CNO administration does not alter sleep amount, architecture or slow-wave activity in a DREADD-free control group (R26^LSL-hM4Di^). **(a)** Schematic of electrophysiological recordings in R26^LSL-hM4Di^ mice (scale bars represent 500 μV on the y-axis and 1 s on the x-axis) and counterbalanced intraperitoneal (i.p.) injections of saline versus 0.5 mg/kg clozapine-N-oxide (CNO) administered at light onset (ZT=0). **(b)** Representative hypnograms and EEG slow-wave activity (SWA, 0.5-4 Hz, 4-s epochs) of a mouse following injections of saline (top panel) and 0.5 mg/kg CNO (bottom panel). **(c)** Time courses of Wake, NREM, and REM sleep amount in the first 6 hr post-i.p. injection of saline (grey, S) and CNO (brown, C), with corresponding quantifications of time spent in each vigilance state during the first hour of DREADD activation. Rapid eye movement (REM) sleep is presented as a percentage of total sleep time (% TST). **(d)** Representative EEG and EMG traces after saline and CNO injections, with brief awakening examples in yellow highlight. **(e)** Time course of brief awakenings (BA; <16 s movement during sleep; number per hour total sleep time, expressed as % saline recording average) during the first 6 hr after saline (S) and CNO (C), with the corresponding number of BA’s in the first hour post-i.p. injection. **(f)** Time course of EEG NREM slow-wave activity (SWA, % saline recording average) for the first 6 hr after saline (S) and CNO (C), with the corresponding % of NREM SWA in the first hour post-i.p. Injection. **(g and h)** Sleep architecture analyses during **(g)** NREM sleep and **(h)** REM sleep. Note that all time analyses were aligned from 15 min post-injection (as T0) to ensure full DREADD activation and are visualised as mean ± SEM (shaded). Asterisks indicate comparisons with significant differences (*p<0.05; **p<0.01; ***p<0.001) for analyses with significant main effects. n=9 for panels c, g and h, n=8 for panels d, e and f. C: CNO, Clozapine-*N*-oxide; DREADD: Designer Receptors Exclusively Activated by Designer Drugs; EEG: Electroencephalogram; EEG F: Frontal EEG; EEG O: Occipital EEG; EMG: Electromyogram; i.p.: intraperitoneal; NREM: Non-rapid eye movement sleep; REM: Rapid eye movement sleep; S: Saline; SWA: Slow-wave activity; TST: Total sleep time; T0: timepoint of analysis onset 15 min post-injection; ZT: Zeitgeber time.

**Suppl. Fig. 2:**
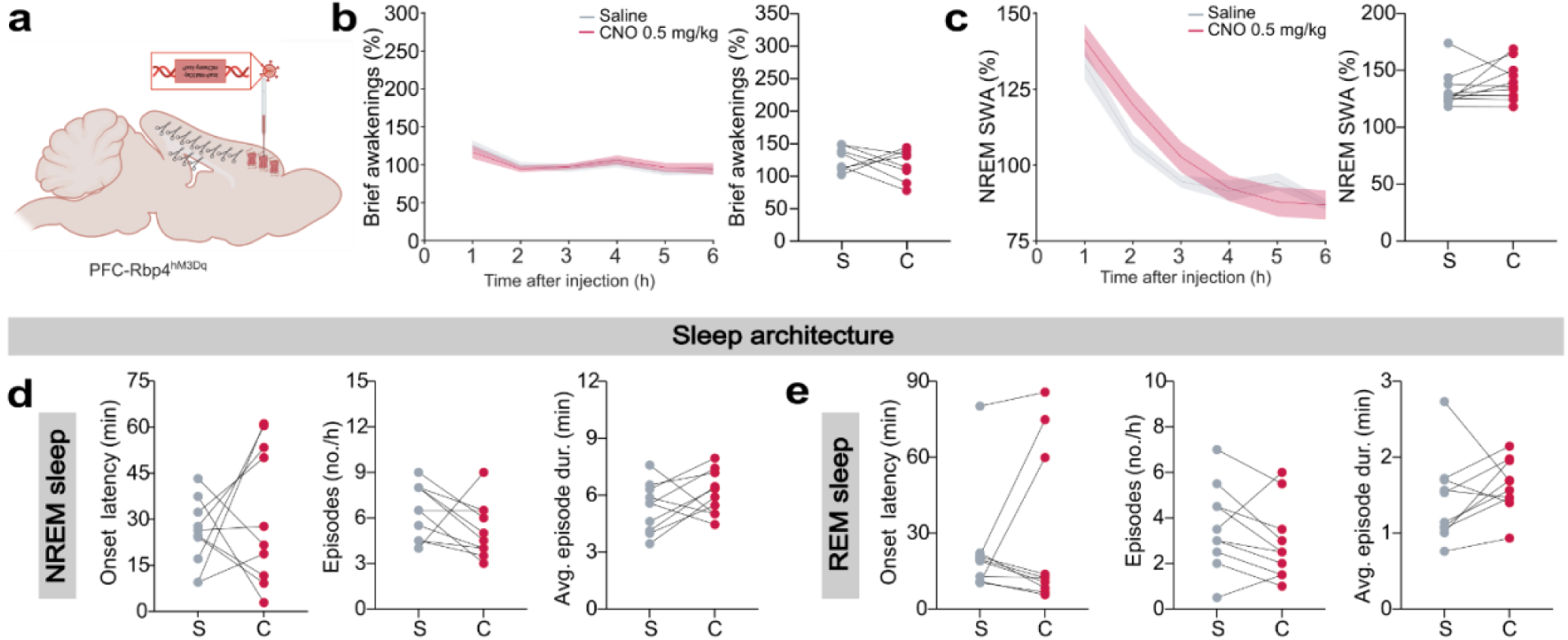
Sleep architecture is unaffected in PFC-Rbp4^hM3Dq^ mice. **(a)** Schematic of viral vector injections for localised hM3Dq DREADD expression in the prefrontal cortex (PFC) of Rpb4^Cre^ mice. Note Cre expression is represented as biological scissors. Mice received counterbalanced intraperitoneal (i.p.) injections of saline and 0.5 mg/kg CNO at light onset (ZT=0), as in Figure 2. **(b)** Time course of brief awakenings (BA; <16 s movement during sleep; number per hour total sleep time, expressed as % saline recording average) during the first 6 hr after saline (S, grey) and CNO (C, red), with the corresponding number of BA’s in the first hour post-i.p. Injection. **(c)** Time course of EEG NREM slow-wave activity (SWA, % saline recording average) for the first 6 hr after saline (S) and CNO (C), with the corresponding % of NREM SWA in the first hour post-i.p. injection. **(d and e)** Sleep architecture analyses during **(d)** NREM sleep and **(e)** REM sleep. Note that all time analyses were aligned from 15 min post-injection (as T0) to ensure full DREADD activation and are visualised as mean ± SEM (shaded). n=8 for panel b, n=10 for panels c, d and e. C: CNO, Clozapine-*N*-oxide; i.p.: intraperitoneal; NREM: Non-rapid eye movement sleep; REM: Rapid eye movement sleep; S: Saline; TST: Total sleep time; T0: timepoint of analysis onset 15 min post-injection; ZT: Zeitgeber time.

**Suppl. Fig. 3:**
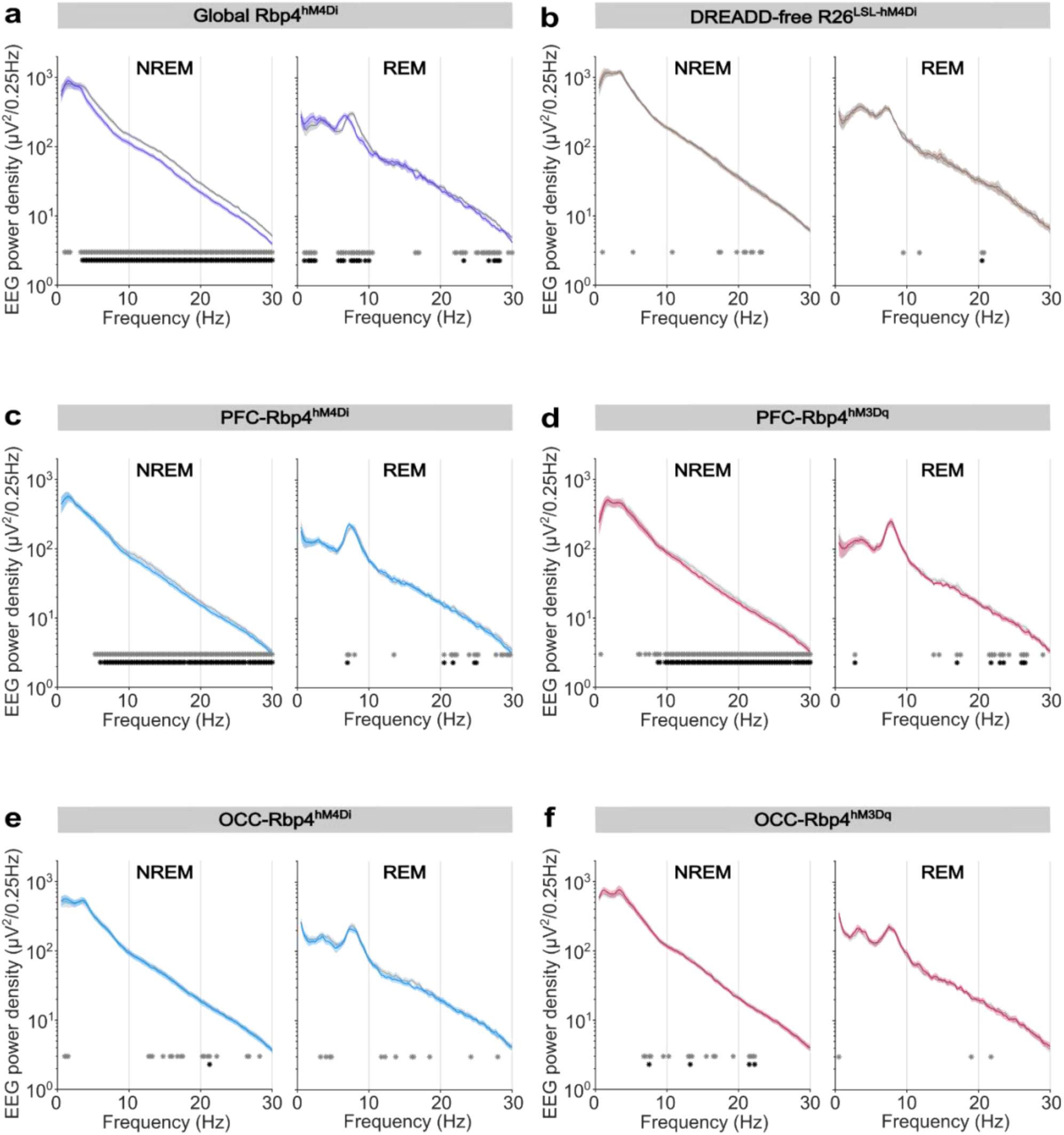
NREM and REM frontal EEG spectra across all experimental groups. Frontal EEG spectra in **(a)** global Rbp4^hM4Di^, **(b)** DREADD-free R26^LSL-hM4Di^, **(c)** PFC-Rbp4^hM4Di^, **(d)** PFC-Rbp4^hM3Dq^, **(e)** OCC-Rbp4^hM4Di^, and **(f)** OCC-Rbp4^hM3Dq^ mice during NREM sleep (left panels) and REM sleep (right panels) in hours 1+2 post-intraperitoneal (i.p.) injection of saline (grey) and 0.5 mg/kg CNO (respective colour) at light onset (ZT=0). Note that all time analyses were aligned from 15 min post-injection (as T0) to ensure full DREADD activation and are visualised as mean ± SEM (shaded). n=12 global Rbp4^hM4Di^ mice, n=8 control R26^LSL-hM4Di^ mice, n=8 PFC-Rbp4^hM4Di^, n=10 PFC-Rbp4^hM3Dq^, n=10 OCC-Rbp4^hM4Di^, n=7 OCC-Rbp4^hM3Dq^ for panels a-f, respectively. Asterisks indicate post-hoc contrasts with significant differences (grey and coloured inlays *p<0.05; black *p<0.01). CNO: Clozapine-*N*-oxide; DREADD: Designer Receptors Exclusively Activated by Designer Drugs; EEG: Electroencephalogram; i.p.: Intraperitoneal; NREM: Non-rapid eye movement sleep; REM: Rapid eye movement sleep; SWA: Slow-wave activity; T0: timepoint of analysis onset 15 min post-injection; ZT: Zeitgeber time.

**Suppl. Video 1: Reduced exploratory behaviour and earlier sleep onset following CNO administration in Rbp4^hM4Di^ mice.**

One-hour time-lapsed video (60x speed) of two representative Rbp4^hM4Di^ mice for 1 hr post-i.p. injection of saline (left mouse) and 5 mg/kg CNO (right mouse) at light onset (ZT=0). Immediately following the i.p. injection, the mice’s own nesting material was scattered evenly across the cage and a novel object (taped toilet roll) was introduced. Camera light sensors were deactivated to generate black and white footage. CNO: Clozapine-N-oxide; i.p.: Intraperitoneal; ZT: Zeitgeber time. Video available at: https://doi.org/10.6084/m9.figshare.30069778

## Notes

https://doi.org/10.6084/m9.figshare.30069778

### Competing Interest Statement

The authors have declared no competing interest.

### Summary of Updates

We have restructured our manuscript to focus on the divergent effects of global and local cortical activity upon sleep. We have also considerably expanded our dataset by bidirectionally modulating activity levels in two different local cortical regions, the prefrontal cortex (PFC) and occipital cortex (OCC), finding that these result in distinct and complementary sleep phenotypes.

